# BOLDsωimsuite: A new software suite for forward modeling of the BOLD fMRI signal

**DOI:** 10.1101/2024.01.28.577679

**Authors:** Jacob Chausse, Avery J. L. Berman, J. Jean Chen

## Abstract

In trying to elucidate the mechanisms of the blood-oxygenation level-dependent (BOLD) functional MRI (fMRI) techniques and to expand on the potential of the transverse relaxation time (T2*) in quantitative MRI, many methods for the forward modelling of the BOLD effect have been created and analyzed. Simulations of this nature can be difficult to implement without prior experience, and differences made by methodological choices can be unclear, which provides a significant barrier of entry into the field. In this paper, we present BOLDswimsuite, a toolbox for forward modeling of the BOLD effect, which collects many of the principal methods used in the literature into a single coherent package. Implemented as a Python package, simulations are built-in scripts by combining various simulation components, while providing flexibility in methodological choices. The goal of this toolbox is to provide an open-source, reproducible software suite that is adaptable for simulations in different MRI applications, and to which additional features can be added by the user with relative ease. This paper first provides an overview of the methods available in the package and how these methods can be constructed from the toolbox’s modular code components. Then, a brief theoretical explanation of each simulation component is given, supported by the relevant contributors. Next, sample simulations and analyses that can be created using the package are presented to display its features. Finally, recommendations regarding computational requirements are included to help users choose the best simulation methods to fit their needs. This package has many use cases and significantly reduces methodological barriers to forward modeling. It can also be a good learning tool for MR physics as well as a powerful tool to promote reproducible science.

## Introduction

While the blood-oxygenation level-dependent (BOLD) fMRI technique has become widely used, the full complexities of its origins remain poorly understood by most users. In addition to studying the effect of blood oxygenation, the transverse relaxation (*T*_2_*)-weighted signal underlying the BOLD technique has also increasingly been used in quantitative MRI applications such as iron quantification (Kor et al., 2019) and the modeling of contrast agent-induced MR signals (Báez-Yánez et al., 2017; Bieri and Scheffler, 2007; Pannetier et al., 2013). Modeling of the *T*_2_*-weighted BOLD signal not only helps to deepen the understanding of BOLD signal origins, but is also invaluable in consolidating the interpretation and signal changes in the above applications.

Forward modeling of the BOLD signal involves the use of microscopic physics to simulate bulk BOLD signals. In the early days after the inception of the BOLD technique, Yablonskiy et al. established physical models of BOLD contrast based on the case of static dephasing (Yablonskiy and Haacke, 1994). Subsequently, making use of the analytical expression of *T*_2_*that links the BOLD signal to perturber geometry (specifically cylinders), (Ogawa et al., 1993), early work demonstrated that the static dephasing regime applies to specific vessel sizes and oxygenation level, and that the influence of diffusion is vessel-size selective (Boxerman et al., 1995a; Kennan et al., 1994). Pioneering the Monte Carlo approach to modeling water diffusion, Boxerman et al. work was especially important for the understanding of microvascular BOLD, which is highly sensitive diffusion and thus to dynamic dephasing. Since then, many approaches to diffusion sensitivity have been introduced. Monte Carlo-based methods have become established as the main-stream method for diffusion modeling in BOLD fMRI (Stone et al., 2019)(Martindale et al., 2008)(Gagnon et al., 2015)(Boxerman et al., 1995a, 1995b). However, a deterministic method instead of the Monte Carlo approach to simulating diffusion was proposed by Bandettini and Wong (Bandettini and Wong, 1995), and has also been successfully applied (Pannetier et al., 2014)(Klassen and Menon, 2007). Amongst these approaches are two-(2D) and three-dimensional (3D) variants, with the 3D approach being more physically realistic but also entailing more computational costs. These different approaches have different advantages and drawbacks, which will be discussed later.

Despite the increasing utility of BOLD forward modeling, the number of research groups familiar with its implementation remains limited, due to the intricate details involved. Open science and reproducible science have received important attention in recent years, and given the various approaches to BOLD forward modeling, and the increasing number of applications for forward modeling of the *T*_2_*-weighted signal decay, it is imperative to enable a more consistent approach to forward modeling across studies. In keeping with this spirit, Pannetier et al. introduced a publicly distributed BOLD modeling toolbox implemented in Matlab, named 2DMrVox (Pannetier et al., 2013). This toolbox implements the 2D deterministic diffusion approach and has found usage in clinical research (Lemasson et al., 2016). Inspired by this prior work and by the principles of Open Science, we propose BOLDsωimsuite, a comprehensive simulation toolbox implemented entirely in Python that permits Monte Carlo or deterministic diffusion-based simulations with a range of geometries for the sources of magnetic field perturbations in 2D and 3D. In the following sections, we detail our implementation, their physical principles, highlight some applications, and make recommendations for users.

## Overview of toolbox

The BOLDsωimsuite Toolbox is entirely implemented in Python (Version 3.10). Its various methods are implemented as combinations of methodological choices which are themselves categorized into four components, namely “geometry”, “B0 offset”, “diffusion” and “signal calculation”. Figure 1 shows the four components and their respective options, as well as valid combinations of options. These include

**Figure 1.**
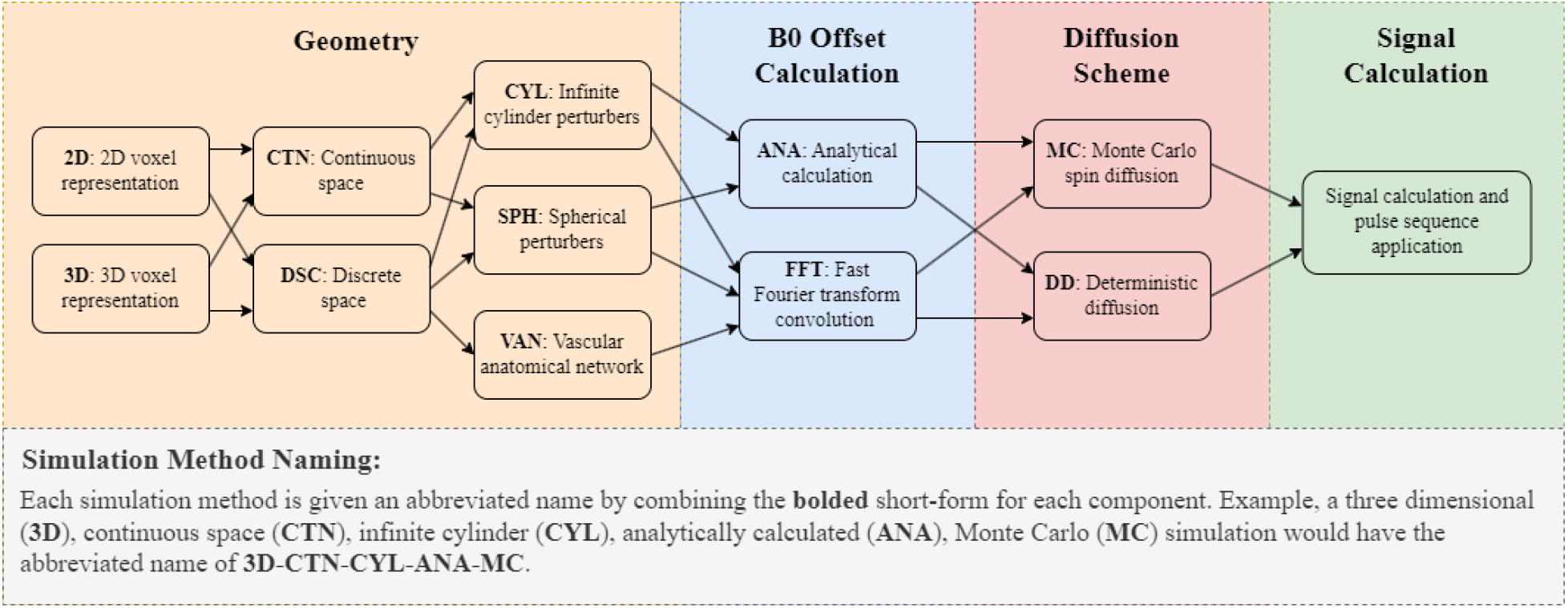
The main components in the numerical simulation pipeline are separated into four categories. Geometry options change the way the vascular network is represented. *B*_0_ offset calculation options change the method used to calculate a perturber’s influence on the magnetic field. Diffusion scheme options depend on the sampling method (discrete or continuous) used for the *B*_0_ offset. Signal calculation contains many parameters that define the pulse sequence and properties of the signal, such as relaxation times.

- Geometry options:
  - Two-vs. three-dimensional (2D vs. 3D);
  - Continuous vs. gridded spatial coordinates;
  - Cylinder, sphere or custom (VAN);
- B0 offset calculation options:
  - Analytical (ANA) vs. Fourier (FFT) field offset calculations;
- Diffusion options:
  - Monte Carlo (MC) diffusion vs. deterministic (DD) diffusion of spins.

These will all be described in detail in the following sections. Most combinations of options can be permuted, with a few exceptions, namely (1) the FFT method can only be applied in discretized space; (2) the custom geometry option can only be used with the FFT method. Representing the toolbox this way allows us to create a short but comprehensive naming scheme which can describe each method. Figure 1 details how these names are generated and how to understand them.

Using the BOLDsωimsuite package, simulations are run in Python scripts built by the user. Figure 2 shows a more in-depth view of the internal structure of the package, including functions and objects involved in a BOLD signal simulation. Depending on the simulation method desired, a specific subset of objects and functions are required to add all the necessary simulation components. Each of these objects or methods have attributes or parameters that need to be defined. These may be simple inputs or could be other objects (which need to be defined themselves). Presenting the toolbox this way provides a better understanding of how each method can be implemented, and can be used as a preliminary guide for generating working simulation scripts.

**Figure 2.**
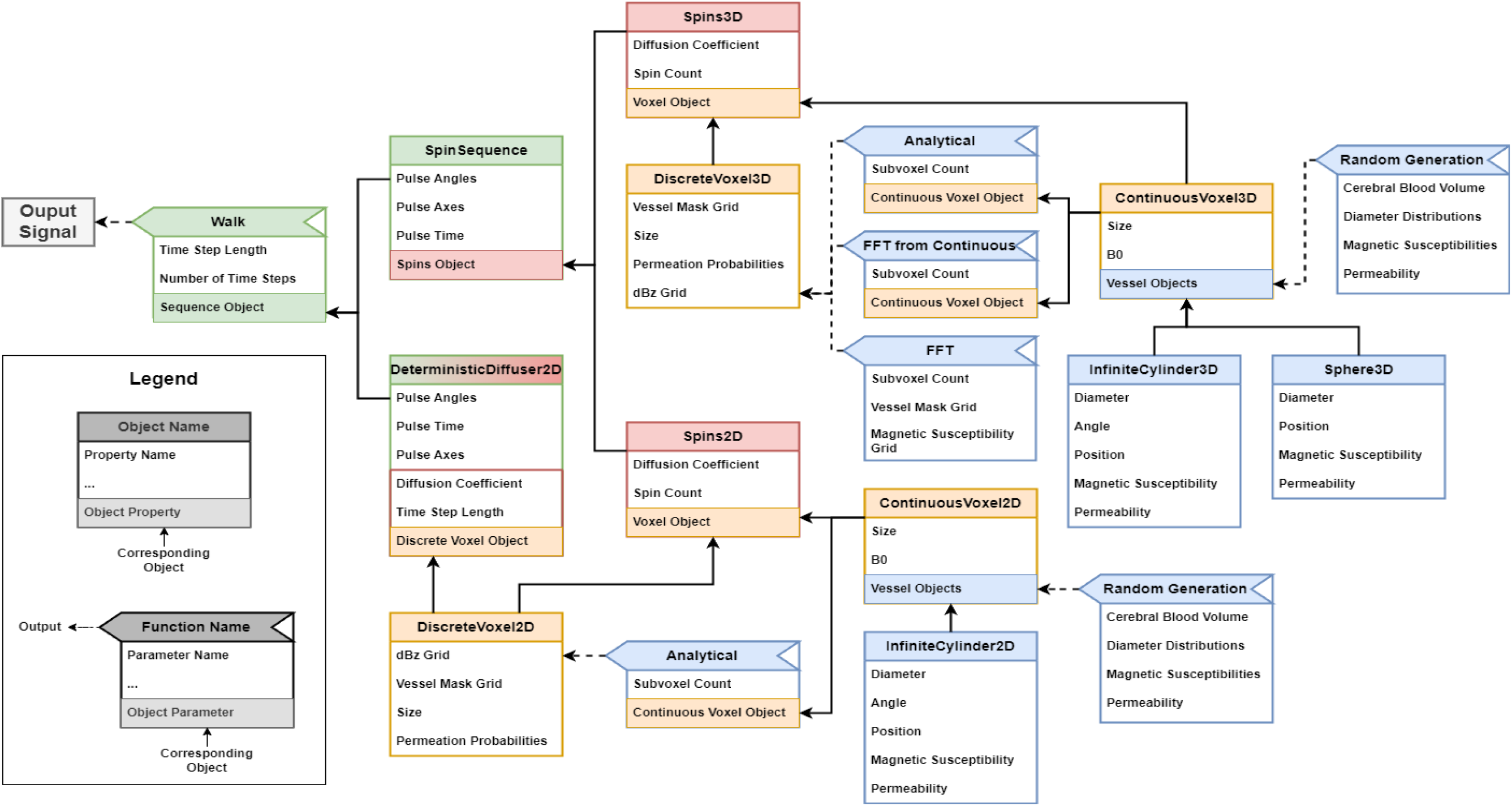
Module flow chart for the BOLDsωimsuite package. All currently possible simulation pipelines are shown for the case of a simple BOLD signal simulation. Each box represents either a class object or a member function, as shown in the legend. In each box are listed the attributes or parameters that are required or can be interacted with. The colour of each object/method relates to the simulation component it represents, as shown in Figure 1. For example, the ContinuousVoxel3D object is yellow, meaning that it relates to the Geometry component, whereas the Spins2D object is red, meaning that it relates to the Diffusion Scheme component. Here, the DeterministcDiffuser2D object is an exception (coloured red and green), as it takes care of both the Diffusion Scheme and Signal Calculation components. A single path connecting the output signal on the left back towards the far right of the diagram can be used to identify the necessary objects and methods to generate the Python script that will run the desired simulation.

### Theory

All symbols and acronyms relevant to the theoretical description are listed in Table 1.

**Table 1.** list of symbols.

| Symbol | Definition |
| --- | --- |
| $B_0$ | Main magnetic field (Tesla) |
| CBV | Cerebral blood volume (fraction of voxel volume) |
| $D$ | Diffusion coefficient ($\text{mm}^2/\text{s}$) |
| $\mathbf{D}$ | Diffusion kernel matrix |
| $\Delta B_z$ | Magnetic field offset |
| $\Delta\chi$ | Susceptibility difference between intra-cylinder (or intra-sphere) and extra-cylinder (or extra-sphere) spaces |
| $\gamma$ | Gyromagnetic ratio |
| $\Delta\omega$ | Magnetic resonance frequency offset (radian) |
| $\Delta t$ | Diffusion step size (in seconds) |
| $k_x, k_y, k_z$ | Kspace coordinates |
| $N$ | Number of subvoxels |
| $N_{\text{spins}}$ | Number of water spins used in Monte Carlo diffusion |
| $N_{\text{steps}}$ | Number of diffusion steps used in Monte Carlo diffusion |
| $\phi$ | Azimuthal angle, angle between the |
| $r$ | Magnitude of the vector perpendicular from the cylinder axis to the proton location |
| $R$ | Vessel radius (mm) |
| $\mathbf{R}$ | MR relaxation matrix |
| $\sigma$ | Standard deviation of Gaussian diffusion kernel |
| $W$ | Width of voxel |
| $\theta$ | Polar angle, also the angle between the $B_0$ and the cylinder axis |

### Geometry

The definition of the simulation geometry requires three choices: number of dimensions, spatial discretization, and perturber geometry.

#### Dimensionality

The choice between 2D (Berman et al., 2018; Miller and Jezzard, 2008; Ogawa et al., 1993; Pannetier et al., 2014) and 3D (Klassen and Menon, 2007; Martindale et al., 2008) simulations have major impacts on the compatible simulation pipeline and the geometry representation.

3D simulations operate in a cubic volume, while 2D simulations operate on a square plane, both called voxels (3D voxels and 2D voxels). A 3D voxel allows for more flexibility in the simulated geometry as it better represents real vasculature. However, 2D simulations are less computationally demanding and can provide equally accurate results to their 3D counterpart when the added flexibility is not required.

#### Continuous and discrete space simulations

Simulations can be performed either on a continuous or discrete space. Continuous-space simulations offer greater spatial accuracy through the use of analytically defined perturbers and efficient sampling of the field offset. They also tend to be less memory intensive since they do not require large arrays that store spatially varying properties of the voxel (e.g., field offsets and relaxation times).

On the other hand, discretizing space allows for much more flexible perturber geometries (e.g., branching vessels) but incurs a tradeoff between spatial accuracy and computational efficiency. Discretized simulations calculate the magnetic field offset over the entire voxel, with equally-spaced sampling, thereby forming grids (with *N* elements on each side, where *N* is defined by the user).

#### Perturber definition and generation

BOLDsωimsuite can simulate three main classes of perturbers: infinite cylinders, spheres, and custom-defined perturber masks. The first two are defined entirely analytically and are generated using very few parameters, as will be described in the next section. They can be either defined by the user or generated randomly using parameters. Currently, the toolbox offers, by default, the functionality of generating randomly oriented cylinders and spheres in continuous space. The custom-defined perturber masks can accommodate any perturber geometry such as vascular anatomical networks (VANs). These require complete discretized masks, which must be generated externally. Sample perturber configurations are shown in Figure 3.

**Figure 3.**
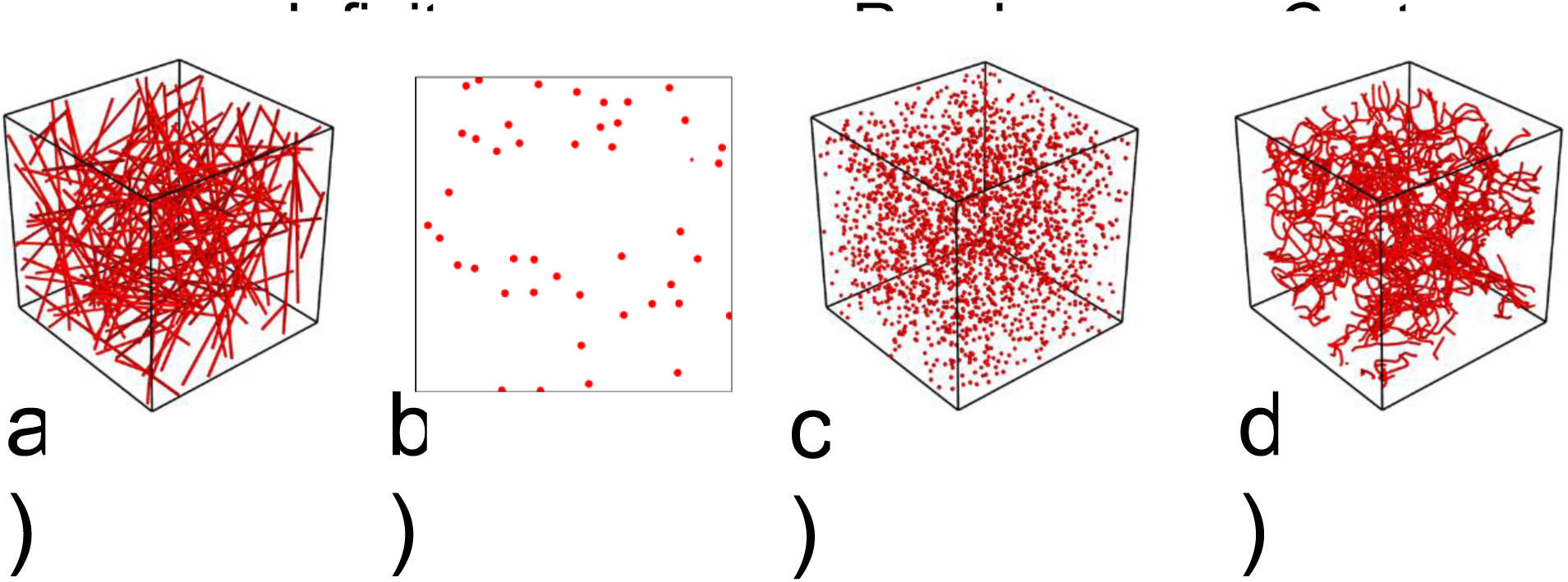
Depictions of sample voxels. Infinite-cylinder models can be constructed in 3D (a) or 2D (b). Microsphere and empirically determined microvascular (VAN) models can be constructed in 3d (c-d, respectively).

#### Infinite Cylinders

Infinite cylinders are a commonly used blood vessel model but are limited by their lack of curvature and branching. As a result, they have a varying degree of accuracy regarding the representation of realistic vasculature.

3D infinite cylinders are defined by their diameter, position, orientation, magnetic susceptibility and water permeation probability (permeability). Both the position and orientation can be randomly generated, while all other parameters must be input manually. The method employed to randomly generate these parameters provides a uniformly distributed CBV (Martindale et al., 2008).

In a 2D simulation, infinite cylinder cross-sections are always taken as the plane perpendicular to the voxel plane. To achieve a field offset distribution that is comparable to randomly oriented 3D infinite cylinders, each cylinder is assigned its own randomly oriented magnetic field direction (Miller and Jezzard, 2008). Each infinite cylinder is defined by its diameter, position, magnetic field orientation, magnetic susceptibility, and permeation probability. Both the position and magnetic field orientation can be randomly generated, while all other parameters must be input manually.

#### Spheres

The spherical perturbers are not meant to represent vasculature, but can instead be used for a variety of other perturbers for more unconventional simulations such as red blood cells, ferritin (Brammerloh et al., 2021), or even polystyrene microspheres used in BOLD phantoms (Bieri and Scheffler, 2007). These perturbers are only implemented for 3D simulations. Spherical perturbers are defined by their diameter, position, magnetic susceptibility, and permeation probability. Like infinite cylinders, their position can be randomly generated while all other parameters must be input manually. Unlike cylinders, however, they do not have an orientation.

#### Custom-defined perturber masks

Custom-defined perturber masks are only available for 3D simulations, and they rely on user-generated perturber geometry. Although this makes them the most flexible perturber, generating the perturber mask usually requires significant work outside the toolbox. For example, VANs can be generated using such methods as 2-photon microscopy (Cheng et al., 2019; Gagnon et al., 2015).

### B0 Field Offset Calculation

The field offset in the simulation can be calculated in two distinct ways, either analytically or by FFT convolution.

#### Analytical

Analytically defined perturbers allow for exact calculation of the magnetic field offset and can be used either in continuous or discrete space simulations. The magnetic field offset of infinite cylinder vessels is defined analytically as follows for the intravascular (IV) and extravascular (EV) spaces, based on (Ogawa et al., 1993),

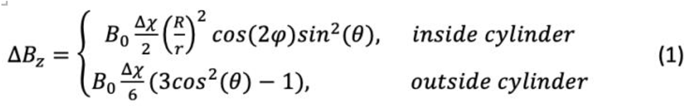

Where in both cases, *B*_0_ is the applied magnetic field strength, Δχ is the magnetic susceptibility difference between the intra- and extravascular spaces. R is the cylinder radius, and *r* represents the distance between the cylinder’s axis and the spin position spanned by the vector **r**, θ represents the angle between the magnetic field direction and the cylinder axis, and ϕ is the angle between the vector **r** and the projection of B0 onto the plane orthogonal to the cylinder axis. The field offset associated with spherical perturbers is modeled as (Boxerman et al., 1995b)

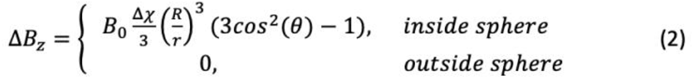

where R is the sphere radius, r represents the distance between the sphere’s origin and the spin position, and θ represents the angle between *B*_0_ and **r**.

#### Fourier convolution

The magnetic field offset of a magnetization distribution can also be calculated through the discrete Fourier transform. The local dipoles of the field perturbations can be calculated in the Fourier domain and then Fourier-transformed back to the spatial domain (Deville et al., 1979). Our implementation is based on the work by Cheng et al. (Cheng et al., 2009), where the 3D B_Z_ is defined in Fourier space as

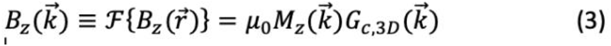

where *B*_z_ is derived by convolution (multiplication in Fourier space) with a kernel *G*,

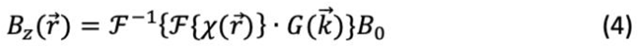

The kernel *G* is first calculated in the spatial domain as,

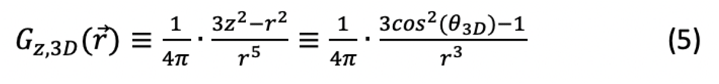

*G*_z_ is set to zero at r = 0, and is subsequently converted to Fourier space as

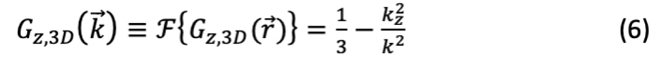

where 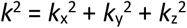, and *G*_c,3D_ is the continuous Fourier transform of the 3D Green’s function. For the 2D case,

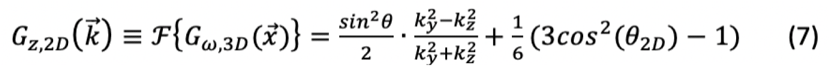

Alternatively,

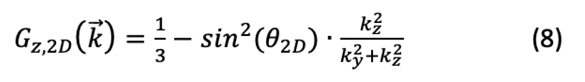

One clear advantage of the Fourier convolution method is its independence from analytical equations. Thus, the perturber layout and shape are arbitrary and can contain any geometry, including in vivo mapped VANs. However, the field offsets generated through Fourier convolutions are inherently discretized and thus have limited spatial resolution. Further, they are susceptible to boundary effects and wrap-around artifacts. To prevent these wrap-around artifacts, it is usually recommended to add zero padding to the arrays used in Fourier transforms (and inverse Fourier transforms). The amount of zero-padding is set by the user, whereby selecting to have no padding will have the effect of mirroring the voxel at its edge boundaries (due to the periodicity of the discrete Fourier representation). This can be desired to emulate the presence of similarly dense vascular networks around the voxel, with the downside that this surrounding is just a mirrored version of the voxel. Setting the zero-padding to half of N will completely remove wrap-around artifacts.

### Diffusion

Simulating diffusion effects using the BOLDsωimsuite can be done in two different ways, either with the gold standard Monte Carlo or using the convolution-based deterministic method.

#### Monte Carlo diffusion

Monte Carlo diffusion uses randomly positioned particles that move in the voxel through a random walk. Since the discovery of the effect of diffusion on BOLD contrast (Ogawa et al., 1993), there have been several efforts to standardize the implementation of Monte Carlo simulations (Boxerman et al., 1995a; Fieremans and Lee, 2018; Martindale et al., 2008), there have been efforts to standardize its implementation (Fieremans and Lee, 2018; Martindale et al., 2008). In our implementation, consistent with these other works, each step length is randomly sampled from a normal distribution with a standard deviation of 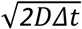, where D is the diffusion coefficient and *Δt* is the time step length. Note, that the particles’ positions are always computed in continuous space, regardless if the simulation geometry is defined in continuous or discrete space. This allows for diffusion steps that are much smaller than the grid elements to cumulatively take effect. Voxel boundary conditions are imposed on the particles to model the behaviour of diffusing water molecules. First, the voxel boundaries are periodic, such that any particle exiting the voxel on one end reappears on the other (and moved inward by the remaining diffusion distance). This is done to ensure that diffusion is not restricted by the voxel boundaries but also that particles do not leave the voxel’s volume.

#### Perturber permeability

Another boundary is formed by the permeability of the pertubers. When a particle step crosses a perturber’s wall, it has a chance to successfully permeate the wall, given by the perturber’s “permeation probability”. If the permeation is successful, the particle moves to the new position, otherwise, a new step is randomly generated until one is found that does not cross the wall. Many simulations use either fully permeable or fully impermeable vessels, but the Monte Carlo diffusion implemented in BOLDsωimsuite allows for partially permeable perturbers.

At each step, the particles accumulate a spin dephasing due to the magnetic field offset. This can be calculated from the B0 offset using the Larmor equation. In the case of continuous space simulations, the dephasing is calculated for each spin at each timestep. For discrete space simulations, the B0 offset is pre-calculated on the entire grid, and spin dephasing is sampled from the grid.

#### Deterministic diffusion

The deterministic diffusion method represents diffusion as a smoothing process of convolving spin magnetizations with a Gaussian kernel, as diffusion of an ensemble of spins is Gaussian distributed (Bandettini and Wong, 1995). The magnetization at time point j is given by

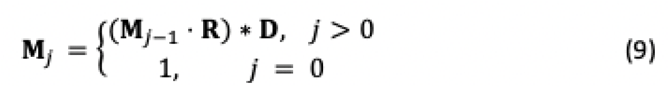

Where j is the time-step index, and **R** represents the *T*2 relaxation process. **D** is the diffusion kernel. We implemented deterministic diffusion for 2D simulations using options of Gaussian and modified Bessel kernels. The Bessel kernel was first introduced (Pannetier et al., 2014) as an improvement over the 2D Gaussian kernel (Lindeberg, n.d.)Elements of the Gaussian diffusion kernel are defined as

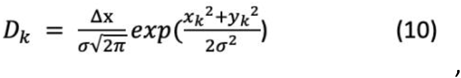

Elements of the Bessel kernel are defined as

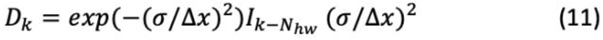

where N_hw_ is the number of samples at half of the kernel width, and *k* is the sample index in the diffusion kernel width direction, and *I*_k_-*N*_hw_ is the modified Bessel function of the first kind (order *k*-*N*_hw_).

Boundary conditions in the convolution are addressed by padding the voxel with itself, such that spins that hit the voxel boundary are wrapped to the other side. This enables the deterministic diffusion process to closely emulate the behaviour of Monte Carlo diffusion.

#### Perturber permeability

In the deterministic diffusion framework, perturbers are defined as fully permeable or impermeable. By default, the kernel convolution does not interact with perturber boundaries, which corresponds to fully permeable perturbers. It is optional to include a correction at each step which emulates impermeable perturber behaviour by effectively returning any magnetization that crossed tissue boundaries back to the original tissue (Pannetier et al., 2014). Currently, no intermediate levels of permeability are implemented.

### Signal calculation

RF pulses are implemented by specifying the pulse axis in polar coordinates. For example, an RF pulse along the x-axis is specified by the orientation vector [π/2, 0], while a pulse along the y-axis is given by [π/2, π/2], where the first angle represents the polar angle, relative to the z-axis, and the second angle represents the azimuthal angle, relative to the +x-direction. Each pulse also requires a flip angle (in radians) and the time step at which it is applied.

At each step, the change in the precessional frequency is determined by

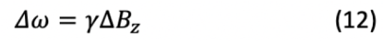

And the additional phase in position p accumulated at each time step due to this frequency shift is given by

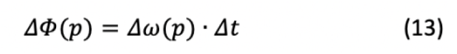

The cumulative phase accumulation over all time steps is given by

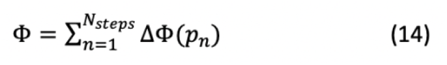

Thus, at each position *p*, the MR signal is defined as

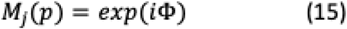

The BOLD signal at each time point j is in turn computed as the complex conjugate of the magnetization at each spin location.

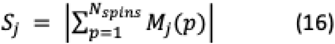

where *N* is the length along each direction of the voxel (number of grid points).

### Example: simulating a spin echo BOLD signal

Shown in Figure 4 are two examples of BOLD signal time courses derived using the 2D Monte Carlo setting of BOLDsωimsuite. Figure 4a illustrates a network of infinite cylinders with random orientations (*θ*), which is typical of cortical grey matter. The resultant dBz map is shown in Figure 4b, and the corresponding BOLD time courses for a spin-echo (SE) sequence (TE = 80 ms) are shown in Figure 4c for the total BOLD signal as well as its intravascular (IV) and extravascular (EV) constituents. We can compute Δχ of fully deoxygenated blood (cgs units) using the relationship given in (Spees et al., 2001). The signal undergoes gradient-echo (GE) *T*_2_* decay up to 40 ms, at which point the refocusing pulse is applied. In Figure 4d we show the spatial map of a population of densely packed cylinders, such as those found in a white matter bundle. Currently, these white matter fibre layouts can be reconstructed in the toolbox from the outputs generated by the AxonPacking package (Mingasson et al., 2017). The circles represent axonal cross sections, and *B*_0_ is assumed to be at a 90° angle to the fibres. This bundle of largely parallel cylinders is interspersed with a few randomly oriented blood vessels. We set Δχ of the axons, also in cgs units, according to past studies (Rudko et al., 2014). Likewise, the spin-echo BOLD signals corresponding to this “white-matter” voxel are shown in Figure 4f.

**Figure 4.**
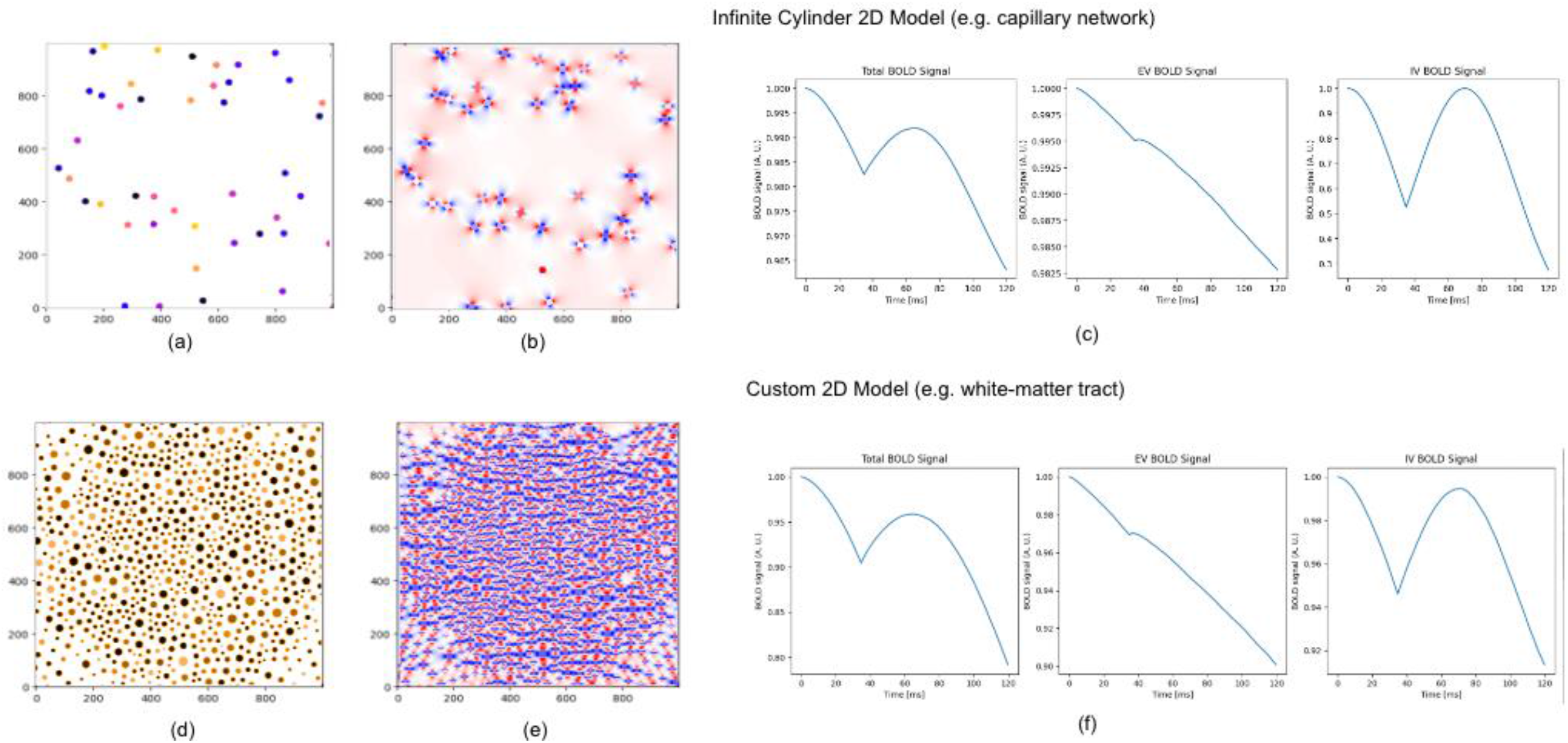
Sample simulated voxel configurations (2D) and corresponding spin-echo BOLD signals. (a) A network of infinite cylinders with random orientations (*θ*) illustrated in a 2D voxel, in which the colours encode different perturbers. (b) The resultant dBz map, in which blue and red indicate the magnetic dipoles surrounding the cylindrical vessels. The axes indicate physical size in arbitrary units. (c) The corresponding BOLD time courses for a spin-echo sequence (TE = 80 ms) for the total BOLD signal as well as its intravascular (IV) and extravascular (EV) constituents. (d) A 2D voxel with densely packed axons, with the circles representing axonal cross sections, and *B*_0_ is assumed to be at a 90° angle to the fibres. (f) The spin-echo BOLD signals corresponding to this “white-matter” voxel.

## Applications

### BOLD signal modeling based on vascular networks

An obvious application for this toolbox is in BOLD signal modeling, an approach that was pioneered by Boxerman et al. (Boxerman et al., 1995b). Boxerman et al. simulated the GE and SE BOLD signals as a function of perturber radius, as well as the IV contribution. As we showed in our previous work (Berman A. J. L., Chausse J., Hartung G., Polimeni J. R., Chen J. J., n.d.), the modeling results derived from different simulation approaches are different. As an example, Figure 5a shows the dependence of the total BOLD signal on the vascular radius, demonstrating broad agreement amongst the 3D-ANA-MC, 2D-ANA-MC and 2D-ANA-DD methods. However, upon closer examination of the IV component, greater differences were found across methods, especially in the case of the GE signal.

**Figure 5.**
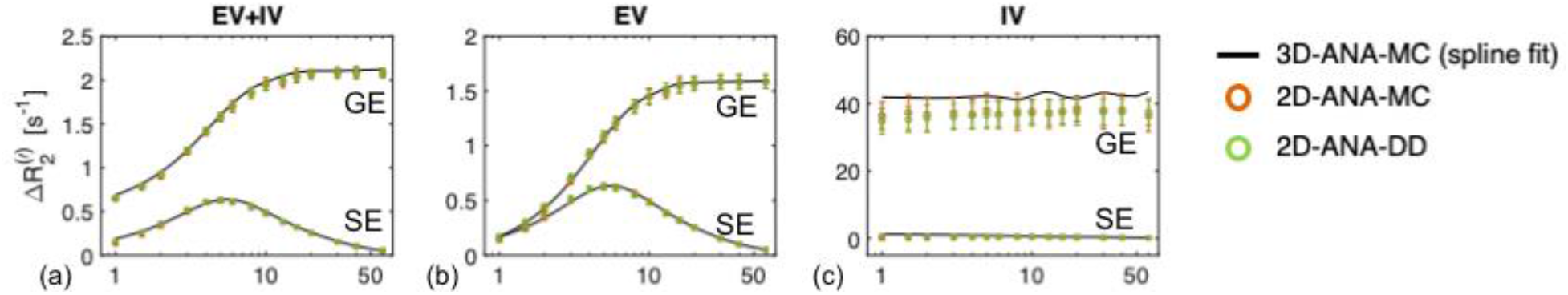
Comparison of Boxerman-style plots showing similarities and differences across simulation approaches. Δ*R*_2_ represents the transverse relaxation attributable to perturbed. Simulations were performed for gradient-(GE) and spin-echo (SE), using 3 different methods, namely 3D-ANA-MC, 2D-ANA-MC and 2D-ANA-DD. For their explanations please refer to Figure 1.

### Quantitative fMRI: calibrated BOLD and vascular MRF

Forward modeling of the *T*_2_ decay due to vascular perturbers was integral to the development of vascular MR fingerprinting (MRvF) techniques (Christen et al., 2014), and is important for the continued development of calibrated BOLD. In the calibrated BOLD model, simulations of the BOLD effect have lent key insight into the microvascular and metabolic contributions to the BOLD signal (Cheng et al., 2019; Gagnon et al., 2015), and continue to enable ways to optimize microvascular (thus neuronal) specificity (Berman et al., 2018). In the MRvF method, estimates of blood oxygenation, volume, and radius are obtained by matching the measured decay to a dictionary of simulated decay curves, the simulated decay curves must be as representative of the experimental conditions as possible. In the original MRvF paper, the 3D infinite-cylinder model with deterministic diffusion was used to generate the dictionary, and contrast enhancement was used to minimize the effect of background macroscopic field inhomogeneities. Christen et al. found different levels of dictionary-matching reliability for the three parameters. Upon close inspection, we found that the dictionary generation process could be influenced by the simulation approach. In our recent work (Berman A. J. L., Chausse J., Hartung G., Polimeni J. R., Chen J. J., n.d.), we demonstrated that using MRvF dictionaries built from different simulation approaches resulted in different error rates in MRvF metrics, as shown in Figure 6. In this case, the ground-truth radius values were used in generating the simulated experimental signal (based on the 3D-ANA-MC) technique. While the use of the same technique in dictionary generation resulted in fully accurate radius estimation results, the use of a dictionary based on the 3D-FFT-MC method resulted in large estimation errors at some radii. Thus, the optimal generation of MRvF dictionaries requires further investigation.

**Figure 6.**
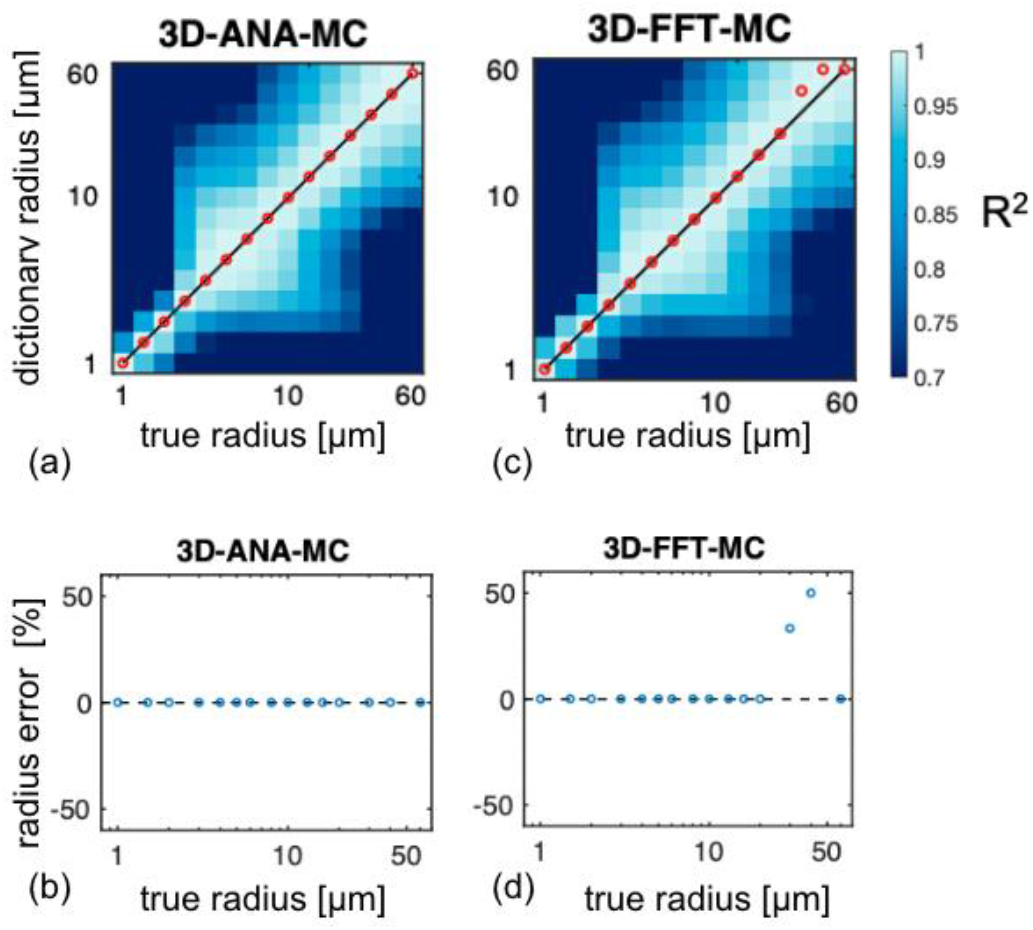
The accuracy of vascular MRF (MRvF) varies by simulation method. The coefficient of variation (R^2^) contour plots (a, c) indicate the uniqueness of the dictionary match at each ground-truth radius. At higher radii, the use of dictionaries generated using the 3D-ANA-MC and 3D-FFT-MC methods resulted in different vessel radius estimates (b, d). Figure adapted from (Berman A. J. L., Chausse J., Hartung G., Polimeni J. R., Chen J. J., n.d.).

### T_2_*-weighted signal modeling of microspheres: red blood cells, contrast agents and iron deposition

Given challenges in constructing tissue phantoms containing microvasculature, microspheres, which are easier to distribute evenly in a phantom medium, were suggested as a surrogate for microvascular networks (Scheffler et al., 2018). Our simulation work (Chausse J., Berman A. J. L., Polimeni J. R., Chen J. J., n.d.), in which we included SE, GE and asymmetric SE (ASE) related transverse relaxation rates (*R*_2_’), where R2’ = 1/T2’, confirmed that microspheres do indeed behave like micro-cylinders in the context of BOLD contrast, as shown in Figure 7.

In in vivo applications, in the work by Boxerman et al. (Boxerman et al., 1995b), to simulate the effect of red blood cell (RBC) movement within a medium containing plasma and the contrast agent, RBCs were modeled as magnetized spherical perturbers. While this work found the flow of RBCs to contribute negligibly to the BOLD signal, the ability to model intravascular magnetized spheres easily extends to magnetized contrast particles (Pannetier et al., 2013). Such applications are not restricted to the brain, where contrast agents remain intravascular, but also to other organs, such as the liver, in which contrast agents can be modeled as uniformly distributed ferrite particles (Hernando et al., 2012; Yablonskiy and Haacke, 1994).

**Figure 7.**
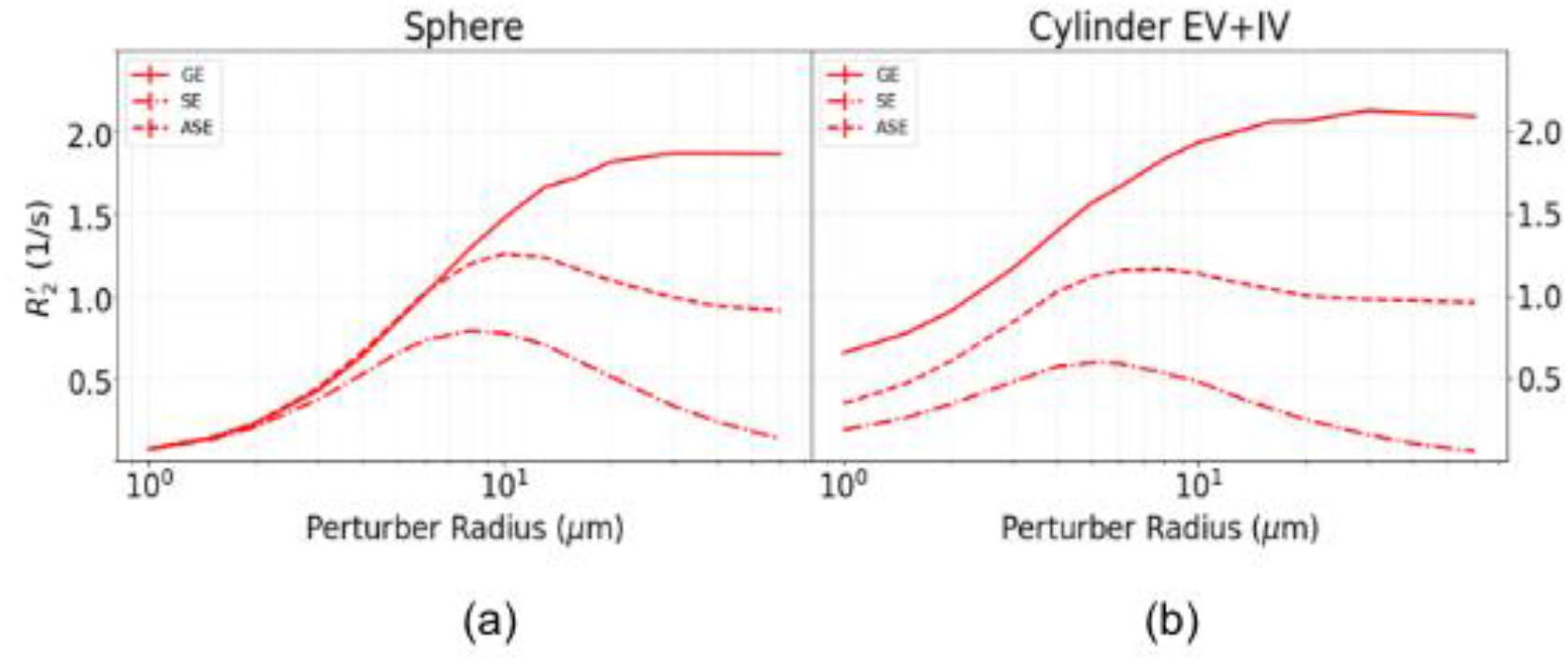
Comparison of Boxerman-style plots for micro-cylinder and microsphere-based voxel models, showing the dependence of transverse relaxation (*R*_2_’) on perturber radius. (a) EV R2’ based on a voxel of microspheres (note that microspheres are assumed to not have IV contributions); (b) the *R*_2_’ plot for infinite cylinders. In both cases, the susceptibility of all perturbed was assumed to be governed by an oxygenation level (Y) of 60%. The perturber radii are plotted on a log scale. Results are shown for gradient-echo (GE), spin-echo (SE) and symmetric spin-echo (ASE) cases. Figure adapted from (Chausse J., Berman A. J. L., Polimeni J. R., Chen J. J., n.d.).

**Figure 7.**
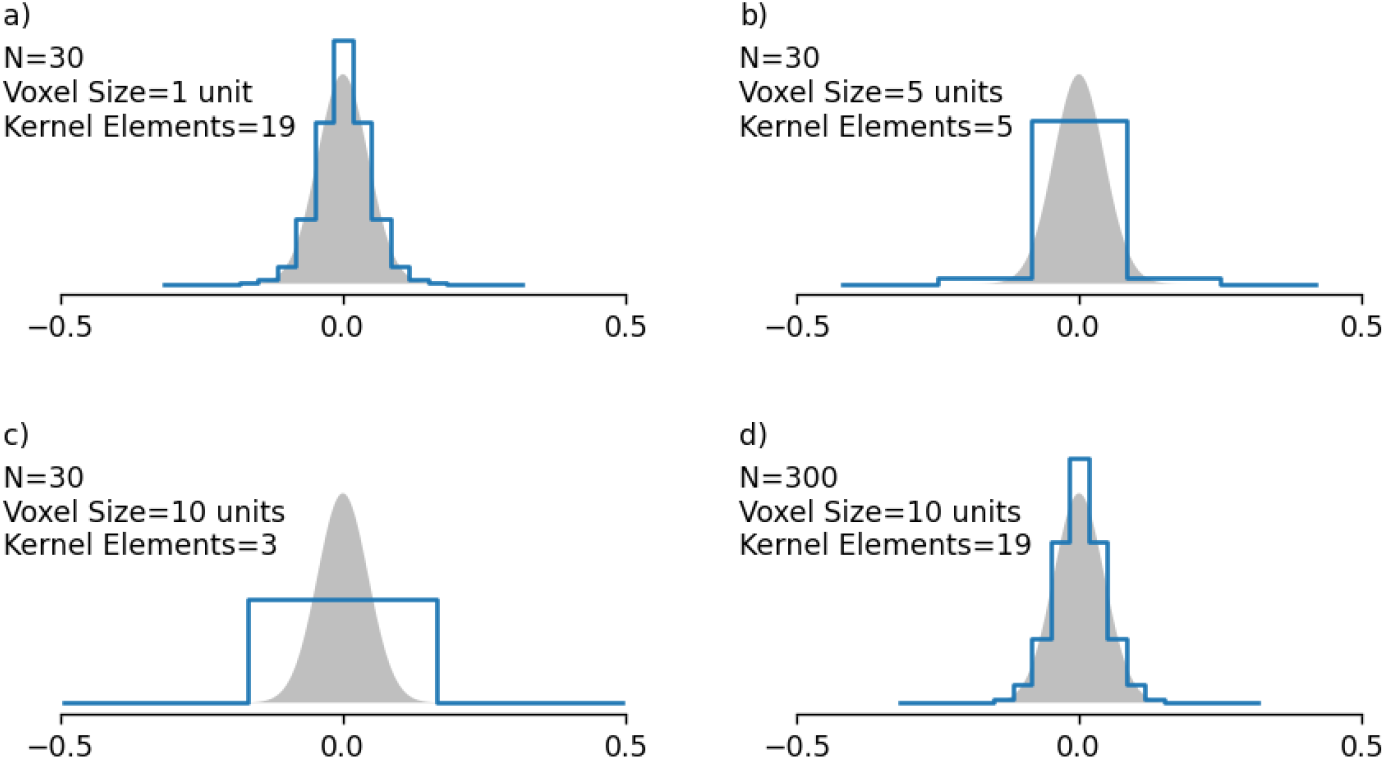
Diffusion kernel (blue line) vs. expected distribution (shaded area), showing over-discretization with increasing voxel size. (a-c) show the diffusion kernel when increasing the voxel size without changing the grid-element count or the diffusion length. This results in a decreasing kernel width. (d) shows the diffusion kernel after the grid-element count in (c) is increased to match the size of each grid element in (a).

In addition to heme iron found in the RBCs, Iron particles are present in vivo in three other main forms, ferritin, transferrin and neuromelanin. Ferritin, for instance, has a spherical shape that is typically about 12 nm in diameter (Mahroum et al., 2022)), and when degraded forms hemosiderin. Ferritin accumulates in the aging process, while hemosiderin concentration is associated with brain disorders and hemorrhage. Different iron molecules induce different magnetic field offsets, and by modeling these molecules as spherical perturbers, one can estimate the concentration of these respective iron species based on T2-weighted MRI or T2 MR relaxometry (Brammerloh et al., 2021).

## Computational considerations

### 2D vs. 3D

In general, 3D simulations are more realistic and versatile. Especially when VANs are concerned, 3D is the only option. We know that 2D vessels can accurately model voxels with randomly oriented vessels (Miller and Jezzard, 2008) and voxels with parallel vessels. Currently, there is no easy way to model arbitrary vascular configurations with 2D simulations.

However, there are certain cases for which a 2D voxel has clear advantages. For instance, the inclusion of randomly oriented blood vessels in densely packed white matter fibres is simplified through the use of a 2D voxel. That is, the vessels and fibres can be modeled as infinite cylinders, whereby random orientations of blood vessels can be accounted for by randomizing the orientation of the effective B0 instead of re-orienting the vessels themselves, which avoids the complexities of orienting blood vessels while avoiding collision and overlap with densely packed surrounding axons.

Moreover, computationally, 2D simulations are much faster than 3D, all simulation parameters being equal. For continuous space simulations, 2D space simplifies the analytical representation of vessels and the movement of spins. Discrete-space simulations also benefit from this, and compared to their 3D counterpart, they have one fewer dimension (of size N) to discretize. Roughly speaking, this speeds up the discretization of space and reduces the memory requirement by a factor of N.

### Continuous vs. Discrete space

For a meaningful comparison, this section only considers cases in which both continuous and discrete space representation are possible. Thus, since custom perturber geometries are only possible with discrete-space simulations, they will not be included in this discussion.

One consideration for discretized simulations is sampling requirements. To allow accurate discrete representation of diffusion and offset effects, the grid must sufficiently sample the perturber, as will also be discussed in the following subsection. We recommend that the smallest perturber be sampled with at least 6 grid elements across its diameter.

In nearly all cases, discrete-space simulations require significantly more memory than continuous space simulations, as the entire voxel space must be discretized and held in memory. On the other hand, for continuous simulations, the voxel’s spatial properties are calculated as needed using analytical equations, resulting in a smaller memory footprint. Because of this, discrete-space 3D simulations should, in most cases, only be used when the discrete-space voxel will be reused multiple times. The discrete spatial properties can be saved to a file, thereby allowing the discretization step to be skipped in successive simulations of the same voxel. Although it still requires much more memory than a continuous-space simulation, any simulations done on the pre-computed discrete voxel will be much faster than the continuous-space equivalent. However, when repetitions are not involved, continuous space will be faster, in addition to being more accurate.

For analytically-defined voxels, discrete-space simulations can use either analytical or FFT methods for the field offset calculation. For all but the simplest voxel geometries, the FFT method will be faster. This is because the analytical method’s computation time increases linearly with the number of analytical perturbers, and its upfront computation time is very low. This makes it very fast for voxels with few perturbers. In the FFT case, the computation time increases linearly as additional perturbers are added, but at a lower rate. However, its upfront computation time is much higher, which means that although it is less efficient for voxels with few perturbers, it rapidly becomes more efficient as more perturbers are added. It is also important to note that the recommended zero-padding in the FFT convolution will likely cause it to have higher memory requirements during the field offset calculation than the analytical method.

### Monte Carlo vs. deterministic

Deterministic diffusion simulations have the advantage of providing the same result every time, given the same parameters. Monte Carlo simulations by nature do not provide the same results unless provided with the same starting point (seed). They also suffer from varying levels of signal noise, which can be difficult to adequately reduce.

The computational requirements of deterministic methods can vary widely, as the voxel grid element count needs to be high enough to accurately represent the water diffusion distribution given by the diffusion kernel. Figure 7 demonstrates this with an example. In Fig. 7a, a diffusion kernel has been created for a N=30 voxel of size 1 unit, resulting in a kernel containing 19 grid elements (represented by the blue line). This reasonably approximates the desired continuous function (shaded area). Fig. 7b and 7c show the same kernel creation process but for voxels of size 5 and 10 units, respectively (N remains the same). This could be the case for simulating voxels containing larger vasculature. We see here that due to the reduced grid resolution, the kernel contains fewer grid elements, no longer representing a reasonable approximation of the desired kernel function. This is problematic as it causes erroneous diffusion behaviour and leads to a lack of diffusion effect in the extreme case. The only solution for this is to increase N to maintain the same grid resolution, as shown in Fig. 7d. However, doing so significantly increases both simulation time and memory requirements. It is important to note that this discretization problem is not nearly as impactful in Monte Carlo simulations, because our Monte Carlo simulations record the positions of diffusing spins in continuous space, irrespective of whether or not the voxel is discretized. As a result, though the field offset and vessel boundaries are discretized, the diffusion behaviour is not.

It is recommended that 2D diffusion simulations for 100% vascular permeability be implemented with deterministic diffusion. This will in most cases be faster than the Monte Carlo equivalent. This is because full vessel permeability adds significant noise to the output signal in Monte Carlo simulations, but deterministic simulations do not have this issue and will thus provide a noiseless signal without any changes to the parameters.

## Conclusions

In this paper, we presented a Python-based software toolbox, referred to as BOLDsωimsuite, that contains a suite of BOLD-signal forward modeling functionalities to suit modern computing hardware and infrastructures. This software suite, due to its comprehensiveness and flexibility, can facilitate reproducible science in the domain of quantitative MRI.

## Acknowledgments

We thank the Canadian Institutes of Health Research (CIHR) for financial support for this work (JJC). We also thank Dr. Grant Hartung for helping to generate vascular anatomical networks that were illustrated in this work.

